# A sunflower FLS2 guides reconstruction of immune-evasive *Xanthomonas* flg22 recognition across divergent FLS2 backgrounds

**DOI:** 10.64898/2026.10.07.757162

**Authors:** Lichi Zhong, Ling Wu, Qiang Cheng

## Abstract

Immune-evasive *Xanthomonas* flg22 epitopes pose a challenge to FLS2-mediated immune surveillance. A further challenge is to reconstruct their recognition across divergent FLS2 backgrounds with a limited number of amino-acid substitutions. AlphaFold 3 (AF3) screening of 256 natural FLS2 homologues identified sunflower HaFLS2-1, responsive to flg22^Xcc^ and flg22^Xoo^ in transient ROS assays. We validated flg22^Xcc^ recognition by *in planta* ligand-association assays and immune responses in stable *Arabidopsis* lines. Guided by HaFLS2-1 ligand-proximal residues, iterative AF3 searches generated PsFLS2 (Salicaceae) and TjFLS2 (Apocynaceae) variants recognising flg22^Xcc^ with ten or fewer substitutions. Single-residue reversions revealed functional contributions from over half of recurrent substitutions in both backgrounds. RcFLS2 (Rosaceae) variants acquired flg22^Xcc^ responsiveness. In tomato SlFLS2 (Solanaceae), three substitutions significantly enhanced flg22^Xcc^ responses, while all three variants carrying two to four substitutions acquired flg22^Xoo^ responsiveness. These findings demonstrate the utility of AF3-guided natural-receptor discovery and iterative engineering for reconstructing and expanding flg22 recognition.

## Introduction

Plants rely on cell-surface pattern-recognition receptors (PRRs) to detect conserved microbial epitopes, whereas pathogens can evolve sequence variation in these epitopes to evade immune surveillance. FLAGELLIN-SENSING 2 (FLS2) recognition of flg22 provides a well-characterised example of this evolutionary tension. Natural FLS2 diversity encompasses recognition specificities for distinct flagellin variants, including epitopes that evade canonical FLS2 receptors (Fürst et al., 2020; Wei et al., 2020; Wu et al., 2022). Yet the recognition of some immune-evasive flg22 variants remains unresolved despite expanding surveys of natural FLS2 diversity.

The structural basis of FLS2–flg22 recognition provides a foundation for structure-guided receptor engineering. The *Arabidopsis* FLS2–flg22–BAK1 complex is structurally well characterised, and AlphaFold 3 enables modelling of homologous receptor–ligand complexes (Sun et al., 2013; Abramson et al., 2024). Building on these advances, recent studies dissected and engineered FLS2 recognition of immune-evasive flg22 variants from *Agrobacterium tumefaciens* (flg22^Agro^) and *Ralstonia solanacearum* (flg22^Rso^). In sequence-similar FLS2 pairs, expanded recognition was reconstructed through substitutions at ligand- or co-receptor-proximal residues, whereas engineering across more divergent FLS2 backgrounds was achieved through larger leucine-rich repeat (LRR) region exchanges (Li et al., 2025; Zhang et al., 2025). Whether recognition can be reconstructed across more divergent FLS2 backgrounds through a limited number of amino-acid substitutions remains relatively unexplored.

The flg22 epitopes from *Xanthomonas campestris* pv. *campestris* (flg22^Xcc^) and *X. oryzae* pv. *oryzae* (flg22^Xoo^) both contain Val at position 14 and evade recognition by several characterised FLS2 receptors (Wang et al., 2015; Li et al., 2025), with this Val14 residue in flg22^Xcc^ shown to be a key determinant of its reduced elicitation of *Arabidopsis* FLS2-dependent immunity (Sun et al., 2006). We therefore asked whether natural FLS2 diversity contains a receptor capable of recognising flg22^Xcc^ and whether its ligand-proximal residue states could guide reconstruction of recognition in divergent FLS2 backgrounds.

## Results

### AF3-guided screening identifies a sunflower FLS2 that recognises flg22^Xcc^

To explore natural FLS2 receptors potentially compatible with flg22^Xcc^, we used AlphaFold 3 to model binary complexes between flg22^Xcc^ and 256 FLS2 homologues assembled from a previously compiled plant FLS2 dataset (Cheng et al., 2020) and the multi-species receptor-like kinase (RLK) collection of Ngou et al. (2024), with sequences provided in Supplementary Table S1A. One receptor, sunflower HaFLS2-1, stood out from the screen, displaying the highest mean interface predicted TM score (ipTM; 0.850 ± 0.010, mean ± s.d.) and the lowest mean Val14–FLS2 contact PAE (predicted aligned error) across contacts between flg22^Xcc^ Val14 and FLS2 residues within 6 Å (2.491 ± 0.090 Å) (Fig. 1a; Supplementary Table S1B). HaFLS2-1 is one of four FLS2-like paralogues annotated in the sunflower XRQ/B reference genome and has a typical FLS2 architecture comprising an N-terminal signal peptide, 28 extracellular leucine-rich repeats (LRRs), a transmembrane region and a cytoplasmic non-RD kinase domain (Supplementary Fig. S1). In the predicted HaFLS2-1–flg22^Xcc^ complex, flg22^Xcc^ adopted an extended conformation along the concave LRR surface, with the overall binding topology and distribution of ligand-proximal residues closely resembling those observed in the *Arabidopsis* FLS2–flg22–BAK1 crystal structure (PDB 4MN8), which contains flg22 from *Pseudomonas aeruginosa* (flg22^Pae^) (Fig. 1b; Supplementary Fig. S1; Supplementary Table S2).

**Figure 1.**
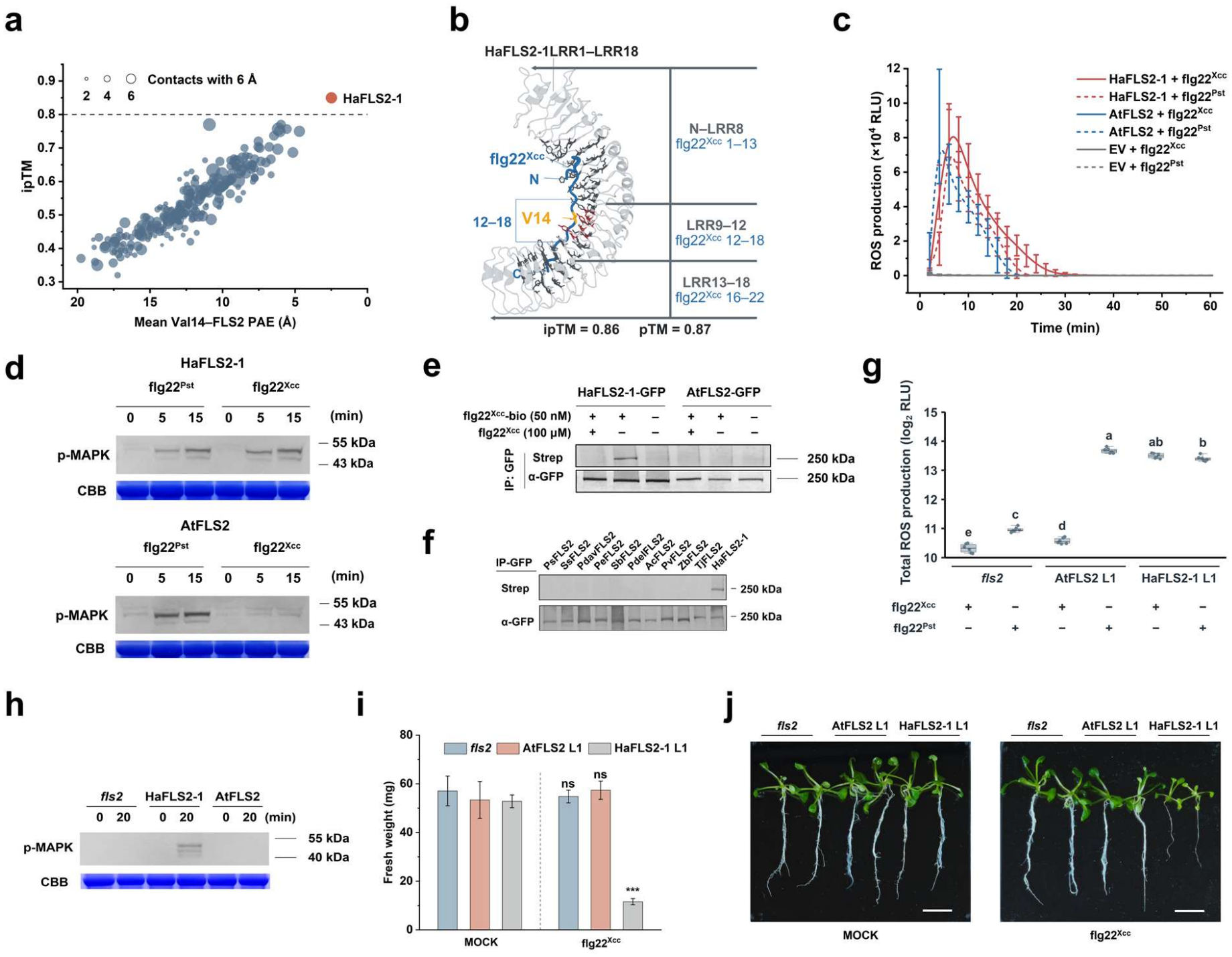
AF3-guided discovery and functional validation of sunflower HaFLS2-1 recognition of flg22^Xcc^. **a**, AlphaFold 3 (AF3) screening of 256 plant FLS2 homologues modelled in binary complexes with flg22^Xcc^. Each point represents one FLS2 homologue. Mean ipTM was calculated across three independent AF3 predictions generated with different random seeds. For each prediction, Val14–FLS2 contact PAE was calculated as the mean bidirectional PAE between flg22^Xcc^ Val14 and FLS2 residues within 6 Å of Val14; the plotted PAE value represents the mean across predictions in which at least one such contact was identified (valid *n* = 1–3). Point size represents the mean number of FLS2 residues within 6 Å of flg22^Xcc^ Val14 across all three predictions. HaFLS2-1 is highlighted. **b**, Representative AF3-predicted model of the HaFLS2-1–flg22^Xcc^ complex. flg22^Xcc^ adopts an extended conformation along the concave surface of HaFLS2-1 spanning LRR1–18. Different portions of flg22^Xcc^ interact with successive regions of the receptor: residues 1–13 with the N-terminal region through LRR8, residues 12–18 with LRR9–12, and residues 16–22 with LRR13–18. flg22^Xcc^ Val14 is highlighted in orange, and ligand-proximal HaFLS2-1 residues are shown as sticks, with Val14-proximal residues highlighted in red. ipTM and pTM values for the displayed model are indicated. **c**, flg22-induced ROS production in *Nicotiana benthamiana fls2* leaves transiently expressing HaFLS2-1 or AtFLS2, or the empty-vector (EV) control. Leaf discs were treated with 200 nM flg22^Xcc^ or flg22^Pst^. Data are mean ± s.d. (*n* = 8 leaf discs per treatment); one representative of three independent infiltration experiments is shown. **d**, MAPK activation in *N. benthamiana fls2* leaves transiently expressing HaFLS2-1 or AtFLS2 following treatment with 1 μM flg22^Pst^ or flg22^Xcc^ for the indicated times. Phosphorylated MAPKs were detected by immunoblotting using a phospho-p44/42 MAPK antibody. Coomassie brilliant blue (CBB) staining is shown as a loading control. One representative of two independent experiments is shown. **e**, Detection of receptor-associated flg22^Xcc^ by an *in planta* ligand-association assay in *N. benthamiana fls2* leaves transiently expressing HaFLS2-1–GFP or AtFLS2–GFP. Leaves were infiltrated with 50 nM C-terminally biotinylated flg22^Xcc^ (flg22^Xcc^-bio) in the presence of 2 mM EGS, with or without 100 μM unlabelled flg22^Xcc^ as a competitor. GFP-tagged receptors were immunoprecipitated using anti-GFP affinity gel. Receptor-associated flg22^Xcc^-bio was detected with alkaline phosphatase-conjugated streptavidin, and immunoprecipitated receptors were detected with an anti-GFP antibody. Signals were visualised by BCIP/NBT staining. **f**, In planta ligand-association assays of FLS2 homologues from *Populus simonii* (PsFLS2), *Salix suchowensis* (SsFLS2), *Populus davidiana* (PdavFLS2), *Populus euphratica* (PeFLS2), *Salix babylonica* (SbFLS2), *Populus deltoides* (PdelFLS2), *Ananas comosus* (AcFLS2), *Phaseolus vulgaris* (PvFLS2), *Zanthoxylum bungeanum* (ZbFLS2) and *Trachelospermum jasminoides* (TjFLS2) in *N. benthamiana fls2* leaves. HaFLS2-1 served as a positive control. Assays were performed as in e. **g**, Total ROS production in stable *Arabidopsis thaliana fls2* transgenic lines expressing AtFLS2 or HaFLS2-1 following treatment with 1 μM flg22^Xcc^ or flg22^Pst^. The *fls2* mutant and representative AtFLS2 and HaFLS2-1 L1 lines were examined. Total ROS production is shown as log^2^-transformed relative light units (RLU). Individual measurements are shown as points, with boxes indicating the median and interquartile range (*n* = 6 leaf discs per treatment). Groups sharing no letter differ significantly (*P* < 0.05). **h**, MAPK activation in the *A. thaliana fls2* mutant and representative L1 transgenic lines expressing HaFLS2-1 or AtFLS2 following treatment with 1 μM flg22^Xcc^ for 0 or 20 min. Phosphorylated MAPKs were detected by immunoblotting, with CBB staining as a loading control, as in d. **i**, flg22^Xcc^-induced seedling growth inhibition in the *A. thaliana fls2* mutant and representative L1 transgenic lines expressing AtFLS2 or HaFLS2-1. Seedlings were grown on solid medium for 7 d and then transferred to liquid medium containing 10 μM flg22^Xcc^ or a DMSO-matched mock medium without peptide. Fresh weight was measured after a further 12 d of growth. Data are mean ± s.d. (*n* = 3 seedlings per genotype and treatment). Comparisons were made between mock and flg22^Xcc^ treatments within each genotype (\*\*\**P* < 0.001; ns, not significant). Corresponding results for the independent L2 transgenic lines are shown in Supplementary Fig. S5. **j**, Representative seedlings corresponding to the growth-inhibition assay in i, showing the *fls2* mutant and AtFLS2 L1 and HaFLS2-1 L1 transgenic lines after mock or 10 μM flg22^Xcc^ treatment for 12 d. Scale bars, 1 cm.

To test whether this predicted structural compatibility translated into functional recognition, we transiently expressed HaFLS2-1 in the *Nicotiana benthamiana fls2* mutant. HaFLS2-1 restored robust reactive oxygen species (ROS) responses to both flg22^Pst^ and flg22^Xcc^, whereas AtFLS2 restored responsiveness only to flg22^Pst^ (Fig. 1c). Consistent with these responses, both peptides induced mitogen-activated protein kinase (MAPK) activation in HaFLS2-1-expressing leaves, while AtFLS2 mediated MAPK activation only in response to flg22^Pst^ (Fig. 1d). Following immunoprecipitation of HaFLS2-1–GFP, receptor-associated biotinylated flg22^Xcc^ was detected using alkaline phosphatase-conjugated streptavidin, and the signal was abolished by excess unlabelled peptide, whereas no corresponding signal was detected for AtFLS2–GFP (Fig. 1e). In addition to AtFLS2, we examined ten FLS2 homologues from different plant species. None exhibited a characteristic flg22^Xcc^-induced ROS burst, although some showed significantly higher total ROS signals than the empty-vector control (Supplementary Fig. S2). In planta ligand-association assays did not detect receptor-associated biotinylated flg22^Xcc^ for any of these ten homologues (Fig. 1f). HaFLS2-1 also responded robustly to flg22^Xoo^ in a broader recognition-spectrum assay (Supplementary Fig. S3). In a parallel AF3 screen of the same 256 FLS2 homologues against flg22^Xoo^, HaFLS2-1 again displayed the highest mean ipTM and lowest mean Val14–FLS2 contact PAE (Supplementary Fig. S4; Supplementary Table S1C).

To further validate HaFLS2-1 function, we generated stable *Arabidopsis fls2* transgenic lines expressing HaFLS2-1 or AtFLS2 under the CaMV 35S promoter. HaFLS2-1 conferred robust flg22^Xcc^-induced ROS production and MAPK activation, whereas AtFLS2 and the *fls2* mutant remained unresponsive (Fig. 1g,h). HaFLS2-1 also conferred flg22^Xcc^-induced seedling growth inhibition in two independent transgenic lines (Fig. 1i,j; Supplementary Fig. S5). Together, these results establish HaFLS2-1 as a functional receptor for flg22^Xcc^ capable of activating downstream immune responses.

### Iterative searches reconstruct and expand flg22 recognition across divergent FLS2 backgrounds

The predicted HaFLS2-1–flg22^Xcc^ complex largely preserved the canonical FLS2–flg22 contact framework (43 of 55 AtFLS2 contact positions), despite substantial divergence in residue identity (19 of 43 identical; Fig. 1b; Supplementary Table S2). To exploit this structural correspondence, we defined ligand-proximal difference positions (LPDs) as ligand-proximal sites identified across predicted HaFLS2-1–flg22^Xcc^ complexes at which a recipient FLS2 differed from HaFLS2-1 (Fig. 2a). We then used HaFLS2-1 residue states at these LPDs to guide iterative searches across divergent FLS2 backgrounds. Candidate variants were generated using three iterative operations—sliding-window (SW) substitution, +1 expansion and −1 reversion—and evaluated using AF3-derived ipTM and Val14–FLS2 contact PAE to prioritise subsequent search steps (Fig. 2b).

**Figure 2.**
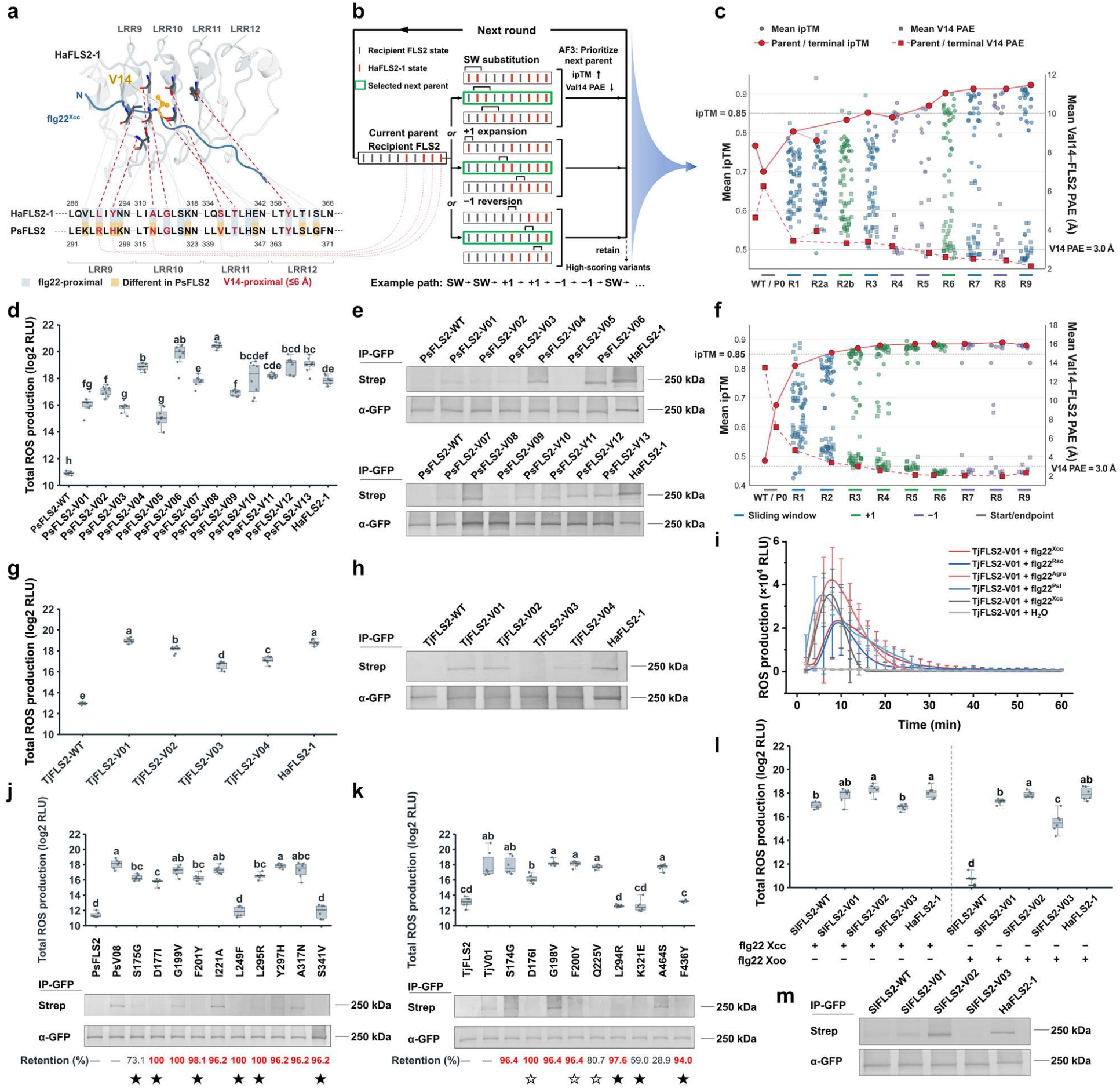
Iterative reconstruction and expansion of flg22 recognition across divergent FLS2 backgrounds. **a**, Structural and sequence comparison used to identify ligand-proximal difference positions (LPDs) and select the starting substitutions for PsFLS2 engineering. A representative AF3-predicted HaFLS2-1–flg22^Xcc^ complex is shown around the Val14-contact region across LRR9–12. Sequence alignment of HaFLS2-1 and Simon poplar (*Populus simonii*) FLS2 (PsFLS2) shows the corresponding LRR concave-face segments, with flg22^Xcc^-proximal positions highlighted in blue and positions differing between the two receptors highlighted in yellow. LPDs were defined as positions where PsFLS2 differed from HaFLS2-1 at flg22^Xcc^-proximal sites identified from AF3-predicted complexes. The four Val14-proximal LPDs (≤6 Å) used to establish the starting PsFLS2 parent (R295L, H297Y, N317A and V341S) are highlighted in red. **b**, Schematic of the iterative AF3-guided search framework. A current parent variant was diversified through three alternative operations: sliding-window (SW) substitution, +1 expansion and −1 reversion. Candidate variants were modelled with flg22^Xcc^, and ipTM and Val14–FLS2 contact PAE were used to prioritise the next parent. A prioritised variant was used as the parent for the next round, while high-scoring variants were retained across rounds. **c**, AF3-guided primary search trajectory for PsFLS2 over nine iterative rounds. Each candidate variant was evaluated using at least three independent AF3 predictions and is represented by a circle (mean ipTM) and a square (mean Val14–FLS2 contact PAE). Mean ipTM was calculated across all predictions, whereas mean Val14–FLS2 contact PAE was calculated across predictions in which at least one FLS2 residue was identified within 6 Å of flg22^Xcc^ Val14. Variants generated by SW substitution, +1 expansion and −1 reversion are distinguished by colour. Red markers and connecting lines indicate the parent or terminal variants selected during the iterative search. Dashed reference lines indicate the empirically defined working thresholds of ipTM = 0.85 and Val14–FLS2 contact PAE = 3.0 Å (see Methods). **d**, flg22^Xcc^-induced ROS production in *Nicotiana benthamiana fls2* leaves transiently expressing wild-type PsFLS2, 13 engineered PsFLS2 variants or HaFLS2-1 (positive control) following treatment with 200 nM flg22^Xcc^. Total ROS production is shown as log^2^-transformed RLU (*n* = 8 leaf discs per treatment). Data presentation and significance notation are as in Fig. 1g. **e**, Detection of receptor-associated flg22^Xcc^ in *N. benthamiana fls2* leaves transiently expressing wild-type PsFLS2, the indicated engineered PsFLS2 variants or HaFLS2-1 (positive control). In planta ligand-association assays were performed as described in Fig. 1e. **f**, AF3-guided primary search trajectory for TjFLS2 over nine iterative rounds. Each candidate variant was evaluated using two independent AF3 predictions. Plotting conventions, calculation of mean ipTM and Val14–FLS2 contact PAE, and empirical working thresholds are as described in c. **g**, flg22^Xcc^-induced ROS production in *N. benthamiana fls2* leaves transiently expressing wild-type TjFLS2, four engineered TjFLS2 variants or HaFLS2-1 (positive control) following treatment with 200 nM flg22^Xcc^. Total ROS production is shown as log_2_-transformed RLU (*n* = 8 leaf discs per treatment). Data presentation and significance notation are as in Fig. 1g. **h**, Detection of receptor-associated flg22^Xcc^ in *N. benthamiana fls2* leaves transiently expressing wild-type TjFLS2, the indicated engineered TjFLS2 variants or HaFLS2-1 (positive control). In planta ligand-association assays were performed as described in Fig. 1e. **i**, Time-course ROS production in *N. benthamiana fls2* leaves transiently expressing TjFLS2-V01 following treatment with 200 nM flg22^Pst^, flg22^Agro^, flg22^Rso^, flg22^Xcc^ or flg22^Xoo^. Water treatment served as the negative control. ROS assays were performed as described in Fig. 1c. Data are mean ± s.d. (*n* = 6 leaf discs per treatment). **j**, Effects of single-residue reversions in PsFLS2-V08 on ROS production induced by 200 nM flg22^Xcc^ (upper) and receptor-associated flg22^Xcc^ signals (lower). Total ROS production is shown as log_2_-transformed RLU (*n* = 6 leaf discs per treatment). Data presentation and significance notation are as in Fig. 1g. In planta ligand-association assays were performed as described in Fig. 1e. Retention percentages indicate the frequencies of the corresponding engineered residue states among the 52 high-scoring PsFLS2 variants from the primary nine-round search (Supplementary Table S6A). Values ≥90% are highlighted in red. Reversion-associated AF3 metrics are provided in Supplementary Table S6B. **k**, Effects of single-residue reversions in TjFLS2-V01 on ROS production induced by 200 nM flg22^Xcc^ (upper) and receptor-associated flg22^Xcc^ signals (lower). Total ROS production is shown as log_2_-transformed RLU (*n* = 6 leaf discs per treatment). Data presentation and significance notation are as in Fig. 1g. In planta ligand-association assays were performed as described in Fig. 1e. Retention percentages indicate the frequencies of the corresponding engineered residue states among the 83 high-scoring TjFLS2 variants from the primary nine-round search (Supplementary Table S6A). Values ≥90% are highlighted in red. Reversion-associated AF3 metrics are provided in Supplementary Table S6B. In j and k, filled stars (★) indicate single-residue reversions that significantly reduced flg22^Xcc^-induced ROS production relative to the corresponding engineered parent and weakened or rendered receptor-associated flg22^Xcc^ signals undetectable. Open stars (☆) indicate reversions that weakened receptor-associated flg22^Xcc^ signals without significantly reducing ROS production. **l**, ROS production in *N. benthamiana fls2* leaves transiently expressing wild-type SlFLS2, three engineered SlFLS2 variants or HaFLS2-1 (positive control) following treatment with 200 nM flg22^Xcc^ or flg22^Xoo^. Total ROS production is shown as log_2_-transformed RLU (*n* = 6 leaf discs per treatment). Data presentation and significance notation are as in Fig. 1g. Statistical comparisons were performed separately for each peptide. Corresponding ROS kinetics are shown in Supplementary Fig. S11. **m**, Detection of receptor-associated flg22^Xcc^ in *N. benthamiana fls2* leaves transiently expressing wild-type SlFLS2, three engineered SlFLS2 variants (V01–V03) or HaFLS2-1 (positive control). In planta ligand-association assays were performed as described in Fig. 1e.

For Simon poplar (*Populus simonii*) FLS2 (PsFLS2), this comparison identified 41 LPDs (Fig. 2a; Supplementary Fig. S6). Four LPDs surrounding flg22^Xcc^ Val14 were substituted with the corresponding HaFLS2-1 residue states (R295L, H297Y, N317A and V341S) to establish the starting parent. Nine rounds of iterative search yielded 52 distinct variants meeting the thresholds of ipTM ≥ 0.85 and Val14–FLS2 contact PAE < 3.0 Å (empirically defined; see Methods; Fig. 2c; Supplementary Table S3). Following additional local scans (Supplementary Table S3D), 13 high-scoring variants carrying 7–15 substitutions and representing different sequence combinations were selected for experimental validation (Supplementary Table S5). All 13 variants exhibited significantly enhanced flg22^Xcc^-induced ROS production relative to wild-type PsFLS2 (Fig. 2d; Supplementary Fig. S8), and 12 showed detectable receptor-associated flg22^Xcc^, whereas wild-type PsFLS2 showed neither response (Fig. 2e). ROS production and receptor-associated ligand signals were broadly concordant across variants but did not consistently follow AF3 metrics: V04 (10 substitutions) showed strong signals in both assays (ipTM 0.887; Val14–FLS2 contact PAE 2.594 Å), whereas V05 (13 substitutions) showed weaker signals despite more favourable metrics (ipTM 0.897; Val14–FLS2 contact PAE 2.189 Å).

Applying the same framework to *Trachelospermum jasminoides* FLS2 (TjFLS2), we identified 40 LPDs relative to HaFLS2-1 (Supplementary Fig. S7). Three LPDs surrounding flg22^Xcc^ Val14 were substituted with the corresponding HaFLS2-1 residue states (R294L, H316A and V340S) to establish the starting parent. Nine rounds of iterative search yielded 83 distinct variants meeting the same AF3 thresholds (Fig. 2f; Supplementary Table S4). Four variants carrying 8–11 substitutions were selected for experimental validation (Supplementary Table S5). All four showed significantly enhanced flg22^Xcc^-induced ROS production relative to wild-type TjFLS2 and detectable receptor-associated flg22^Xcc^, whereas wild-type TjFLS2 showed neither response (Fig. 2g,h; Supplementary Fig. S8). V01 (nine substitutions) showed strong ROS and receptor-associated ligand signals alongside favourable AF3 metrics (ipTM 0.890; Val14–FLS2 contact PAE 1.967 Å). However, V03 exhibited weaker signals in both assays despite AF3 metrics comparable to V02 and more favourable than V04.

We next examined the recognition spectra of PsFLS2-V08 and TjFLS2-V01 against five flg22 variants. PsFLS2-V08 responded to flg22^Pst^ and flg22^Xcc^ but not to flg22^Agro^, flg22^Rso^ or flg22^Xoo^ (Supplementary Fig. S9). In contrast, TjFLS2-V01 retained responsiveness to flg22^Pst^, flg22^Agro^ and flg22^Rso^ while gaining recognition of flg22^Xcc^ and flg22^Xoo^, thus responding to all five variants tested (Fig. 2i; Supplementary Fig. S9).

Among the 52 high-scoring PsFLS2 and 83 high-scoring TjFLS2 variants, nine and six substitutions, respectively, were retained in at least 90% of sequences (Supplementary Table S6A). To assess their functional contributions, we performed single-residue reversions in PsFLS2-V08 and TjFLS2-V01. Five of the nine recurrent substitutions in PsFLS2 and four of the six in TjFLS2 contributed to flg22^Xcc^-induced ROS production and/or receptor-associated ligand signals (Fig. 2j–k; Supplementary Table S6B). Less frequently retained substitutions also contributed, including PsFLS2 G175S (73.1%) and TjFLS2 V225Q (80.7%) and E321K (59.0%). Notably, PsFLS2 revertants G199V (ipTM 0.800; Val14–FLS2 contact PAE 3.731 Å), I221A (0.790; 3.211 Å) and A317N (0.847; 3.483 Å) fell outside the AF3 selection criteria while retaining detectable ligand association. Conversely, PsFLS2 S175G (0.870; 2.819 Å) and TjFLS2 K321E (0.875; 2.721 Å) met both thresholds despite weaker or undetectable receptor-associated ligand signals. Thus, AF3 metrics were useful for candidate prioritisation but did not consistently predict the experimental effects of individual substitutions.

We further extended the iterative search to *Rosa chinensis* FLS2 (RcFLS2). Three high-scoring variants carrying 10–11 amino-acid substitutions were constructed and experimentally tested, all of which acquired flg22^Xcc^-induced ROS responsiveness, although receptor-associated flg22^Xcc^ was not detected under the assay conditions (Supplementary Fig. S10; Supplementary Tables S5 and S7).

We next applied the iterative search to tomato (*Solanum lycopersicum*) FLS2 (SlFLS2). Following six consecutive rounds of −1 reversion, we identified a high-scoring variant carrying only two amino-acid substitutions (SlFLS2-V01; mean ipTM 0.915; mean Val14–FLS2 contact PAE 2.857 Å; Supplementary Tables S5 and S8). Three high-scoring variants identified through the iterative search—V01, V02 and V03, carrying two, three and four substitutions, respectively—were selected for experimental validation. Although wild-type SlFLS2 already responded to flg22^Xcc^, V01 and V02 showed earlier ROS peaks, and V02, carrying three amino-acid substitutions, exhibited significantly enhanced total ROS production (Fig. 2l; Supplementary Fig. S11). In planta ligand-association assays revealed a weak but detectable flg22^Xcc^-associated signal for wild-type SlFLS2, whereas V01 and V02 showed stronger signals, with the most pronounced increase observed for V02 (Fig. 2m). Notably, wild-type SlFLS2 showed no detectable ROS response to flg22^Xoo^, whereas all three engineered variants exhibited clear ROS bursts and significantly increased total ROS production (Fig. 2l; Supplementary Fig. S11). Although selected through flg22^Xcc^-guided searches, none of these variants met the empirical AF3 selection thresholds when modelled with flg22^Xoo^ (mean ipTM 0.605–0.700; mean Val14–FLS2 contact PAE 6.527–9.997 Å; Supplementary Tables S8D and S8E).

Together, these results indicate that, although AF3 metrics did not consistently predict experimental responses, they were useful for HaFLS2-1-guided iterative searches that enriched for FLS2 variants responsive to *Xanthomonas* flg22 escape epitopes across divergent receptor backgrounds.

## Discussion

We identified HaFLS2-1 as a natural FLS2 receptor responsive to both flg22^Xcc^ and flg22^Xoo^. Using HaFLS2-1 as a natural sequence template, we conducted relatively small-scale AF3-guided iterative local searches across four recipient FLS2 backgrounds (approximately 970 distinct receptor sequences, 2,600 AF3 predictions and 23 experimentally tested variants). These searches reconstructed flg22^Xcc^ responsiveness in PsFLS2 (Salicaceae), TjFLS2 (Apocynaceae) and RcFLS2 (Rosaceae), and expanded SlFLS2 (Solanaceae) responsiveness to flg22^Xoo^, with representative functional variants carrying 2–10 amino-acid substitutions. These findings suggest potential applications in targeted genome editing of crop *FLS2* genes to enhance recognition of immune-evasive *Xanthomonas* flg22 epitopes.

## Materials and methods

### Plant materials and growth conditions

The previously generated *Nicotiana benthamiana fls2* double mutant was used for transient receptor-expression assays (Wu et al., 2022). Plants were maintained in a controlled growth chamber at 24°C under a 16-h light/8-h dark photoperiod and approximately 60% relative humidity. Four-week-old plants were used for transient receptor expression and subsequent ROS, MAPK and *in planta* ligand-association assays.

The *Arabidopsis thaliana fls2* mutant SALK_062054 in the Columbia (Col-0) background was used for stable transformation. Plants were grown at 22°C under a 16-h light/8-h dark photoperiod.

### FLS2 sequence collection and AF3 structural analysis

FLS2 sequences were assembled from two sources. One set was obtained from the previously compiled plant FLS2 dataset of Cheng et al. (2020). For the second set, protein sequences from the multi-species plant receptor-like kinase (RLK) collection of Ngou et al. (2024) were searched by BLASTP using *A. thaliana* FLS2 (AtFLS2) as the query. Candidate homologues were further examined by reciprocal BLASTP against the *A. thaliana* proteome to verify FLS2 identity. Together, these sequences yielded a panel of 256 FLS2 homologues used for structural screening (Supplementary Table S1A).

Binary complexes of FLS2 extracellular domains with flg22^Xcc^ or flg22^Xoo^ were modelled using AlphaFold 3 (AF3; Abramson et al., 2024). The top-ranked model (model 0) from each independent prediction was used for metric extraction throughout the structural analyses. The interface predicted TM score (ipTM) was obtained directly from the AF3 outputs. To assess local structural confidence around Val14 of each flg22 peptide, FLS2 residues within 6 Å of Val14 were identified using the minimum heavy-atom distance. For each contacting FLS2 residue, the predicted aligned error (PAE) values in both directions, from the FLS2 residue to Val14 and from Val14 to the FLS2 residue, were averaged. These bidirectional PAE values were then averaged across all FLS2 residues within 6 Å of Val14. This locally averaged metric was defined as the Val14–FLS2 contact PAE. If no FLS2 residue was found within 6 Å of Val14, the Val14–FLS2 contact PAE was considered undefined. The number of unique FLS2 residues within 6 Å of Val14 was defined as the Val14 contact-residue count. The Val14– FLS2 contact PAE and Val14 contact-residue count were calculated using a custom script.

For the natural FLS2 screen, three independent AF3 predictions were generated for each receptor–ligand pair using different random seeds. The mean and sample standard deviation (s.d.) of ipTM, Val14–FLS2 contact PAE and Val14 contact-residue count were calculated across independent predictions. Standard deviations were not calculated when only one valid value was available. Individual prediction values, summary statistics and numbers of valid PAE predictions are provided in Supplementary Tables S1B (flg22^Xcc^) and S1C (flg22^Xoo^).

To compare ligand-contact positions between AtFLS2 and HaFLS2-1, receptor residues within 6 Å of any flg22 residue were identified in the AtFLS2–flg22^Pae^ crystal structure (PDB 4MN8) and in the AF3-predicted HaFLS2-1–flg22^Xcc^ complexes using the minimum heavy-atom distance. For HaFLS2-1, only contact positions consistently identified in model 0 of all three independent AF3 predictions were retained for this comparison. Corresponding positions were mapped by sequence alignment, and shared contact positions and their amino-acid identities were determined (Supplementary Table S2).

### Iterative FLS2 engineering and residue-reversion analysis

HaFLS2-1 was used as the natural sequence and structural template for engineering PsFLS2, TjFLS2, RcFLS2 and SlFLS2. First, HaFLS2-1 residues within 6 Å of flg22^Xcc^ were collected across AF3-predicted complexes and deduplicated. HaFLS2-1 was then aligned with each recipient FLS2, and positions at which the recipient amino acids differed from HaFLS2-1 at these ligand-proximal sites were defined as ligand-proximal difference positions (LPDs). This procedure identified 41, 40, 46 and 40 LPDs for PsFLS2, TjFLS2, RcFLS2 and SlFLS2, respectively. At these positions, the corresponding HaFLS2-1 amino acids served as candidate replacement states.

Candidate variants were generated through three operations: sliding-window (SW) substitution, +1 expansion and −1 reversion. For SW scans, candidate LPDs were ordered according to their positions in the recipient FLS2 sequence. Windows encompassing two or three adjacent entries in this ordered list were shifted by one LPD at a time, with the residues within each window replaced by their HaFLS2-1 counterparts. +1 scans introduced one additional HaFLS2-1 residue state, whereas −1 scans restored one engineered position to its original recipient residue state. Candidate sequences were modelled with flg22^Xcc^ using AF3, and mean ipTM and mean Val14– FLS2 contact PAE were jointly considered when selecting parents for subsequent rounds. The ipTM cutoff of 0.85 was based on functionally validated HaFLS2-1, which had a mean ipTM of 0.850. The Val14–FLS2 contact PAE cutoff of 3.0 Å was informed by PsFLS2-V01, which exhibited flg22^Xcc^-induced ROS production and detectable receptor-associated ligand signals despite a mean PAE of 2.948 Å. These values were therefore adopted as empirical working thresholds, with variants showing mean ipTM ≥ 0.85 and mean Val14–FLS2 contact PAE < 3.0 Å classified as high-scoring.

PsFLS2 and TjFLS2 searches were initiated from variants carrying four and three Val14-proximal substitutions, respectively (PsFLS2: R295L, H297Y, N317A and V341S; TjFLS2: R294L, H316A and V340S). Both searches proceeded through nine iterative rounds using SW, +1 and −1 operations. Candidates in the primary PsFLS2 search were evaluated using at least three independent AF3 predictions, whereas TjFLS2 candidates were evaluated using two. Replicate numbers for additional PsFLS2 derivatives varied. The complete search trajectories and replicate-level metrics are provided in Supplementary Tables S3 and S4.

RcFLS2 engineering began with a three-substitution parent (R294L, H316A and V340S). Seven iterative rounds were performed: two rounds of two-position SW substitution, followed by four rounds of single-position addition and a final round of single-position reversion. SlFLS2 engineering similarly began with three substitutions (R296L, H318A and V342S). An initial two-position SW scan was followed by two rounds of single-position addition and six rounds of successive single-position reversion, yielding a two-substitution variant carrying N178D and R296L. Additional variants were examined through a subsequent +1 scan from a three-substitution SlFLS2 parent. RcFLS2 and SlFLS2 engineered candidates were evaluated using two independent AF3 predictions per variant. The RcFLS2 and SlFLS2 search records are provided in Supplementary Tables S7 and S8, respectively.

Selected high-scoring variants of PsFLS2, TjFLS2, RcFLS2 and SlFLS2, identified using the above criteria with flg22^Xcc^, were additionally modelled with flg22^Xoo^ using AF3 to assess their predicted interactions with this related epitope. Throughout the iterative engineering process, flg22^Xcc^ remained the primary ligand used to guide sequence optimisation.

To analyse substitution retention, 52 and 83 unique high-scoring variants from the primary nine-round PsFLS2 and TjFLS2 searches, respectively, were examined. For each engineered position, retention frequency was calculated as the percentage of high-scoring variants carrying the corresponding HaFLS2-1-derived residue state. Residue states retained in at least 90% of variants were classified as recurrent.

To assess the functional contribution of individual substitutions, single-residue revertants of PsFLS2-V08 and TjFLS2-V01 were generated by restoring engineered positions to their corresponding wild-type recipient residues. These revertants were analysed using AF3 and examined for flg22^Xcc^-induced ROS production and *in planta* ligand association. Substitution retention frequencies and reversion-associated AF3 metrics are provided in Supplementary Tables S6A and S6B, respectively.

### Plasmid construction

Coding sequences of the FLS2 receptors and engineered variants used in this study were assembled from synthetic DNA fragments and PCR-amplified fragments. For variants differing by only a small number of substitutions from an existing construct, the required nucleotide changes were introduced through mutagenic primers during PCR amplification. DNA fragments were assembled into the corresponding plant expression vectors by homology-based seamless assembly using the ClonExpress Ultra One Step Cloning Kit V3 (Vazyme, C117). Untagged receptor constructs were used for stable *Arabidopsis* transformation, whereas C-terminally GFP-tagged constructs were used for transient expression in *N. benthamiana* and *in planta* ligand-association assays. All final constructs were verified by Sanger sequencing before use. The amino-acid substitutions, DNA templates, synthetic fragments and primer pairs used to generate the experimentally tested PsFLS2, TjFLS2, RcFLS2 and SlFLS2 variants are provided in Supplementary Table S5.

### *Agrobacterium*-mediated transient expression

FLS2 constructs were introduced into *Agrobacterium tumefaciens* strain GV3101. Bacterial suspensions were prepared in liquid MS0 medium and adjusted to an OD^600^ of 0.2. The suspensions were infiltrated into fully expanded leaves of 4-week-old *N. benthamiana fls2* plants using a needleless syringe. For ROS burst and MAPK activation assays, leaf discs were excised from infiltrated leaves and incubated overnight in sterile water, with peptide treatments initiated at 36 h after infiltration. In planta ligand-association assays were performed at 48 h after infiltration.

### Peptide synthesis

The flg22^Pst^ (TRLSSGLKINSAKDDAAGLQIA), flg22^Rso^ (QRLSTGLRVNSAQDDSAAYAAS), flg22^Agro^ (DRISSGLKVGSASDNAAYWSIA), flg22^Xoo^ (QQLSSGKRITSFAVDAAGGAIA) and flg22^Xcc^ (QQLSSGKRITSASVDAAGLAIS) peptides were synthesised by GenScript at >85% purity. For *in planta* ligand-association assays, flg22^Xcc^ carrying a C-terminal EDA-biotin modification was synthesised by GenScript at the same purity. All peptides were dissolved in dimethyl sulfoxide (DMSO) to prepare 1 mM stock solutions and stored at −20°C until use.

### ROS burst and MAPK activation assays

ROS production was measured using a luminol-based assay. Leaf discs excised from the indicated *N. benthamiana* or *A. thaliana* plants were floated overnight in sterile water in 96-well plates. The water was then replaced with assay solution containing 100 μM luminol, 20 μg/mL horseradish peroxidase and the indicated flg22 peptide at a final concentration of 200 nM for transient expression assays in *N. benthamiana* or 1 μM for stable transgenic *A. thaliana* lines. Luminescence was recorded immediately after peptide addition using a GloMax 96 Microplate Luminometer (Promega). Total ROS production was calculated by integrating relative light units (RLU) over the recording period and expressed as log^2^-transformed total RLU values. Six to eight leaf discs were analysed per treatment, and the experiments were independently repeated three times.

For MAPK activation assays, leaf discs from the indicated *N. benthamiana* or *A. thaliana* plants were floated overnight in sterile water and subsequently treated with 1 μM of the indicated flg22 peptide. Samples were collected at the indicated time points and immediately frozen in liquid nitrogen. Total proteins were extracted using a commercial protein extraction kit (Beyotime) and subjected to immunoblot analysis. Phosphorylated MAPKs were detected using an anti-phospho-p44/42 MAPK (Erk1/2) antibody (Thr202/Tyr204; Cell Signaling Technology). Coomassie brilliant blue staining was used as a loading control. MAPK activation assays were independently repeated twice.

### Generation of stable *Arabidopsis* transgenic lines

Untagged HaFLS2-1 and AtFLS2 constructs driven by the CaMV 35S promoter were introduced into the *A. thaliana fls2* mutant SALK_062054 by *Agrobacterium*-mediated floral-dip transformation. Independent primary transformants were selected using hygromycin resistance and self-pollinated. T2 plants were subsequently screened by hygromycin resistance and PCR analysis. Two independent T2 lines (L1 and L2) were obtained for each receptor construct. Representative L1 lines were used for ROS burst and MAPK activation assays, whereas both independent lines (L1 and L2) were analysed in seedling growth-inhibition assays.

### Seedling growth inhibition assay

Seeds of the indicated *A. thaliana* lines were surface-sterilised and germinated on solid Murashige and Skoog (MS) medium. After 7 d of growth, seedlings were transferred to liquid MS medium containing 10 μM flg22^Xcc^ or to liquid MS medium containing an equivalent concentration of DMSO without peptide as the mock control. After a further 12 d of growth, seedlings were photographed, and the fresh weight of each seedling was measured individually. Both independent transgenic lines (L1 and L2) were examined for each receptor construct.

### In planta ligand-association assay

Receptor–ligand association was examined using an *in planta* cross-linking and immunoprecipitation assay adapted from our previous study (Xiao et al., 2026). At 48 h after *Agrobacterium* infiltration, *N. benthamiana fls2* leaves transiently expressing C-terminally GFP-tagged FLS2 receptors were infiltrated with 50 nM C-terminally biotinylated flg22^Xcc^ (flg22^Xcc^-EDA-biotin) in HEPES buffer (pH 7.5) containing 2 mM ethylene glycol bis(succinimidyl succinate) (EGS; Sangon Biotech, C100221). For competition assays, 100 μM unlabeled flg22^Xcc^ was included together with the biotinylated peptide. Control leaves were infiltrated with the same buffer containing EGS but without peptide. The final DMSO concentration was kept identical across all treatments within each assay. Leaves were incubated at room temperature for 20 min to allow *in planta* cross-linking.

Leaf discs were collected, and total proteins were extracted using Plant RIPA Lysis Buffer (Strong; Beyotime, P0045). GFP-tagged receptors were immunoprecipitated using Anti-GFP Affinity Gel (MedChemExpress, HY-K0229). Immunoprecipitated proteins were separated by 12% SDS-PAGE and transferred onto Hybond ECL membranes. Receptor-associated biotinylated flg22^Xcc^ was detected using alkaline phosphatase-conjugated streptavidin (Sangon Biotech, B110054), whereas immunoprecipitated GFP-tagged receptors were detected using THE™ GFP Antibody, pAb, Rabbit (GenScript, A01704), followed by an alkaline phosphatase-conjugated anti-rabbit IgG secondary antibody. Both signals were visualised using BCIP/NBT substrate solution.

### Statistical analysis

Multiple-group comparisons of log_2_-transformed total ROS production were performed using one-way ANOVA followed by Tukey’s honestly significant difference (HSD) test (Figs. 1g and 2g) or Welch’s one-way ANOVA followed by Games–Howell post hoc tests (Fig. 2d,j–l). Comparisons in Fig. 2l were performed separately for each flg22 peptide. Comparisons between mock and flg22^Xcc^ treatments within each genotype in the seedling growth-inhibition assays were performed using two-tailed unpaired Student’s t-tests. Groups sharing no letter differ significantly at *P* < 0.05. Sample sizes and data presentation are specified in the corresponding figure legends.

## Data availability

The data supporting the findings of this study are provided within the article, its Supplementary Information and accompanying Source Data files. Supplementary Table S1 provides the FLS2 sequences and AF3-derived metrics for the natural receptor screens. Supplementary Tables S2–S8 provide structural contact comparisons, iterative search records, construct information and residue-reversion data. Additional AF3 model outputs are available from the corresponding author upon reasonable request.

## Code availability

Custom scripts used to extract and analyse AF3-derived structural metrics are available from the corresponding author upon reasonable request.

## Author contributions

LZ performed most of the experiments. LZ and QC analysed the data. LW contributed to the experiments. QC conceived the study and wrote the manuscript.

