## Supplementary Figures and Tables for "A sunflower FLS2 guides reconstruction of immune-evasive *Xanthomonas* flg22 recognition across divergent FLS2 backgrounds": Supplementary Figures S1-S11.docx

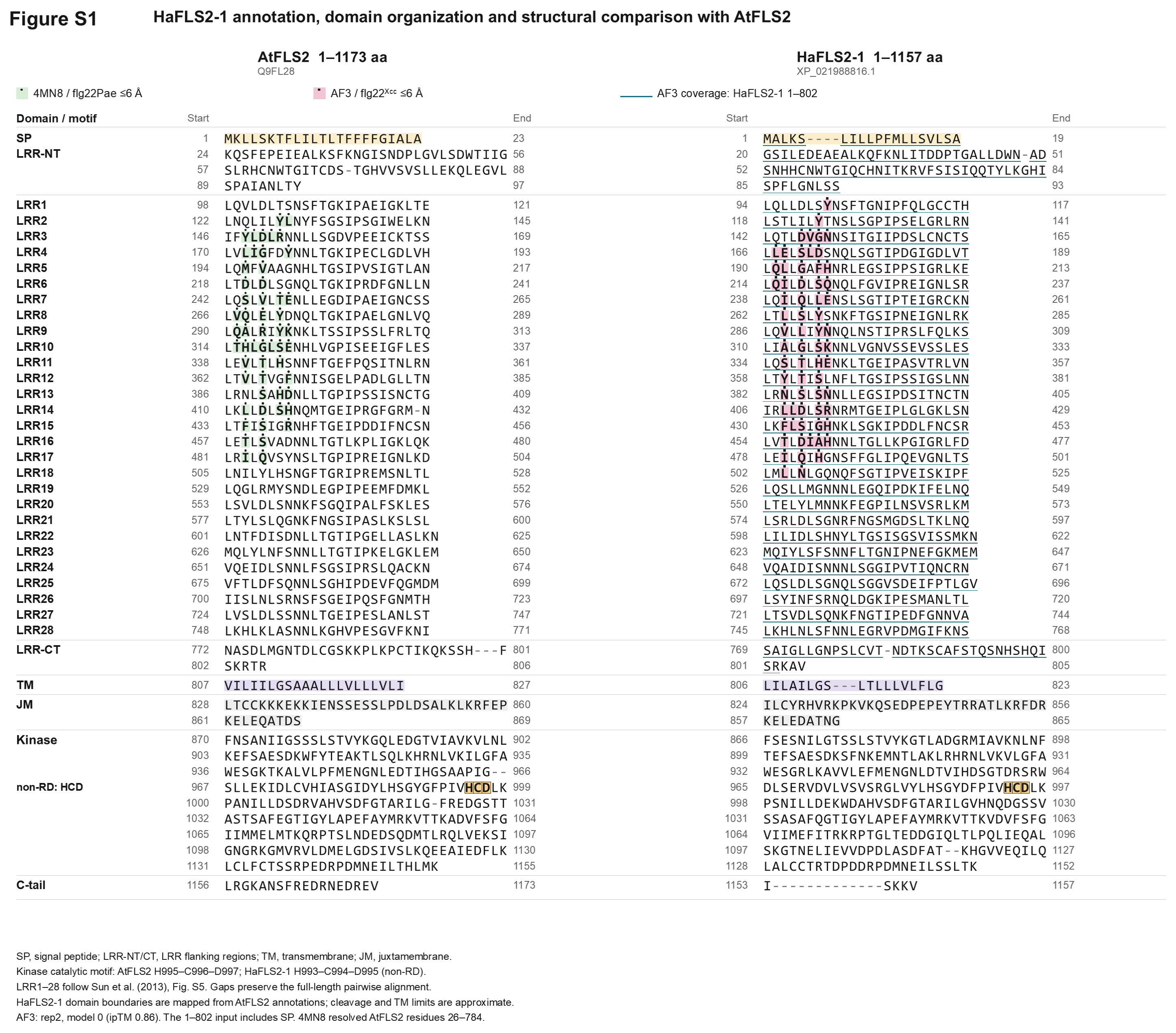

**Figure S1 | Full-length sequence alignment and domain organization of AtFLS2 and HaFLS2-1.** AtFLS2 (UniProt Q9FL28; 1,173 amino acids) and HaFLS2-1 (NCBI XP_021988816.1; 1,157 amino acids) are aligned and organised by domain and leucine-rich repeat (LRR). Residues are numbered from the initiating methionine of each receptor; hyphens indicate alignment gaps. SP, signal peptide; LRR-NT and LRR-CT, N- and C-terminal LRR-flanking regions; TM, transmembrane region; JM, juxtamembrane region; C-tail, C-terminal tail. The HCD motif characteristic of the non-RD kinase domain is indicated. Green shading marks AtFLS2 residues within 6 Å of flg22^Pae^ in the crystal structure PDB 4MN8; pink shading marks ligand-proximal HaFLS2-1 residues in the representative AF3-predicted HaFLS2-1–flg22^Xcc^ model (replicate 2, model 0; ipTM = 0.86). Blue underlining marks the HaFLS2-1 sequence included in the AF3 input (residues 1–802, including the signal peptide). LRR1–28 boundaries follow the AtFLS2 annotation of Sun et al. (2013), with HaFLS2-1 boundaries projected through the alignment. Signal-peptide cleavage and TM boundaries are approximate.

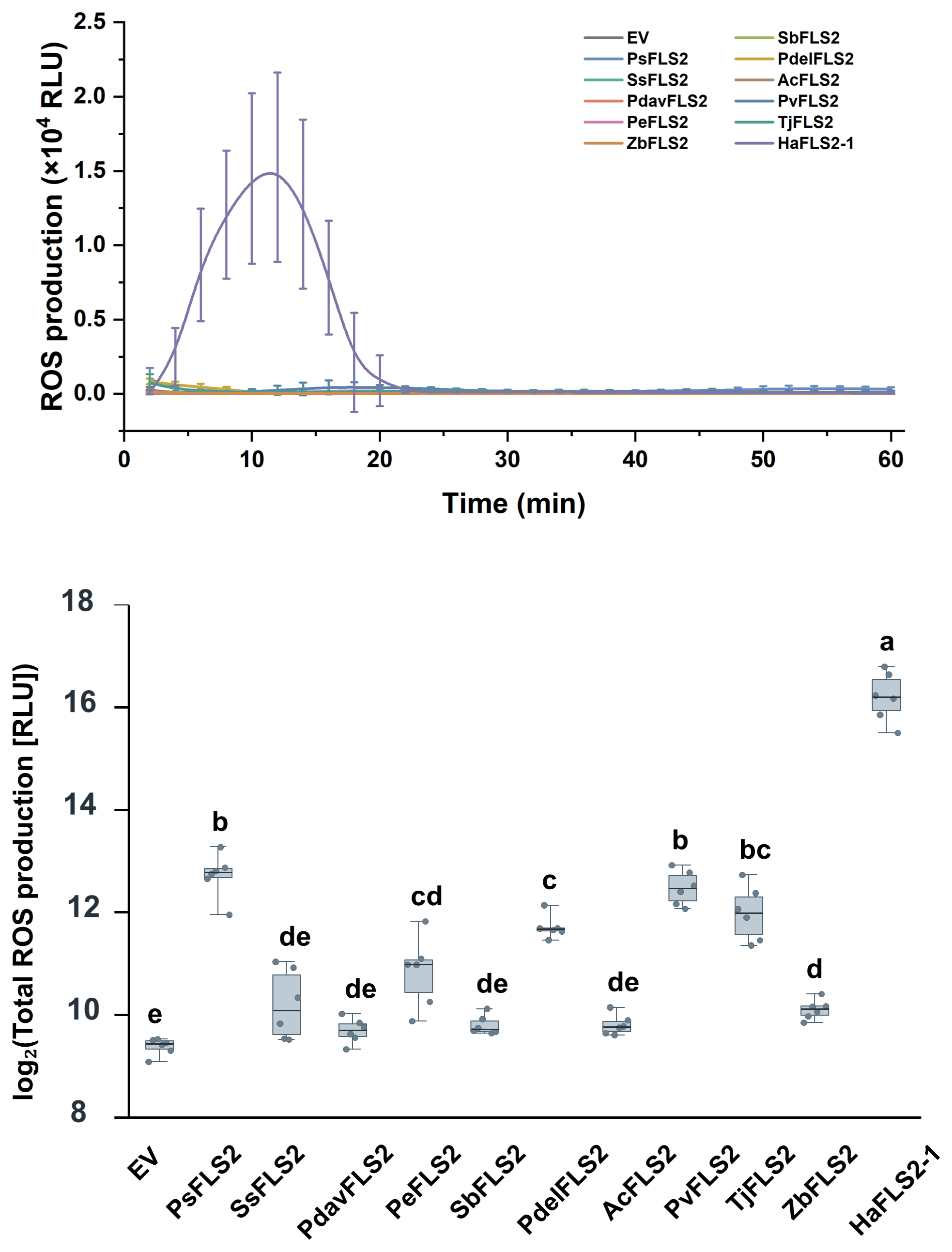

**Figure S2 | flg22^Xcc^-induced ROS production across ten natural FLS2 homologues.** ROS production in *Nicotiana benthamiana* *fls2* leaves transiently expressing FLS2 homologues from *Populus simonii* (PsFLS2), *Salix suchowensis* (SsFLS2), *Populus davidiana* (PdavFLS2), *Populus euphratica* (PeFLS2), *Salix babylonica* (SbFLS2), *Populus deltoides* (PdelFLS2), *Ananas comosus* (AcFLS2), *Phaseolus vulgaris* (PvFLS2), *Trachelospermum jasminoides* (TjFLS2) or *Zanthoxylum bungeanum* (ZbFLS2), following treatment with 200 nM flg22^Xcc^. HaFLS2-1 and the empty vector (EV) served as positive and negative controls, respectively. Top, ROS kinetics over 60 min; curves show mean ± s.d. Bottom, total ROS production shown as log_2_-transformed relative light units (RLU). Individual measurements are shown as points, with boxes indicating the median and interquartile range (*n* = 6 leaf discs per construct). Statistical comparisons used Welch’s one-way ANOVA followed by Games–Howell post hoc tests. Groups sharing no letter differ significantly (*P* < 0.05).

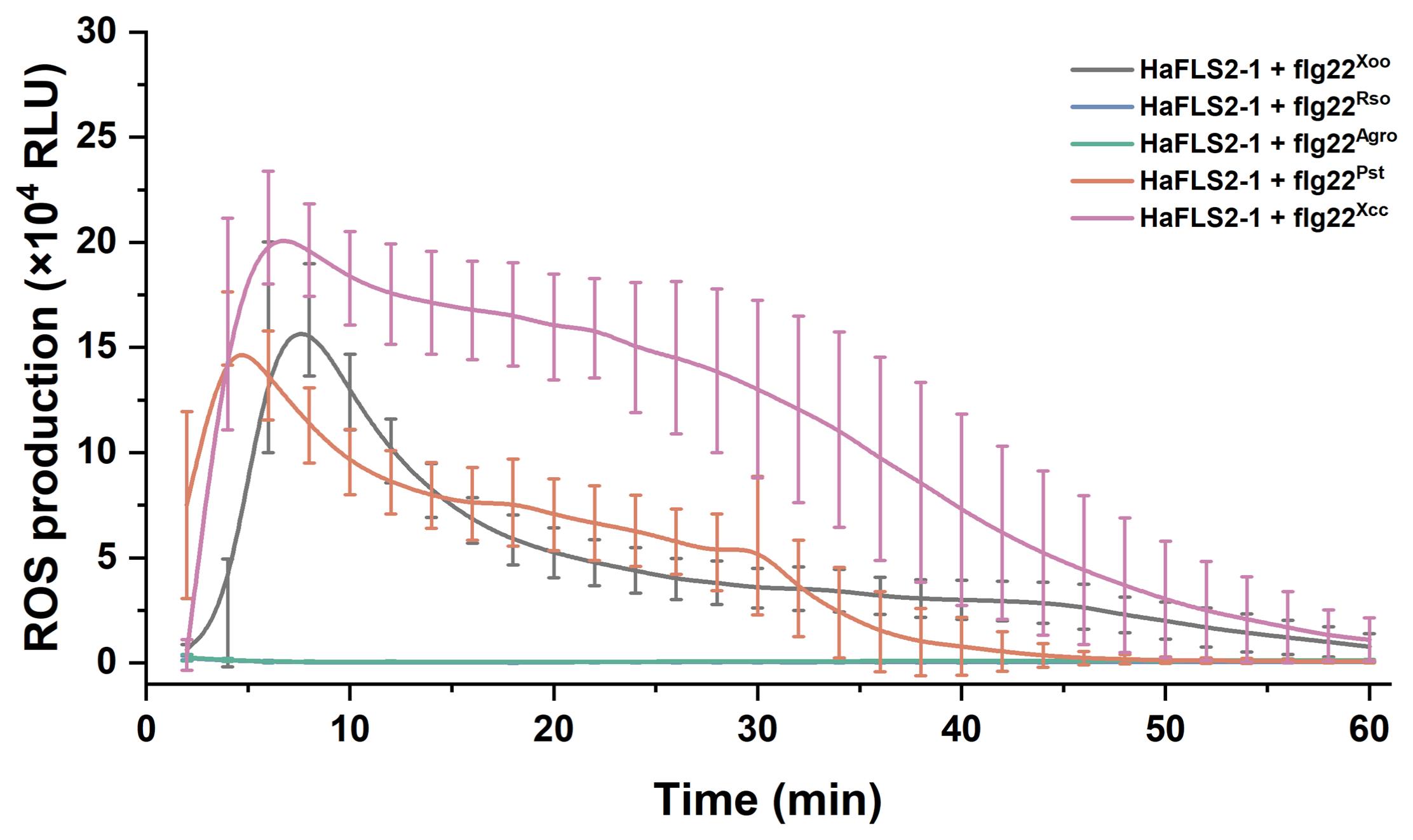

**Figure S3 | Recognition spectrum of HaFLS2-1 across five flg22 variants.** Time-course ROS production in *N. benthamiana* *fls2* leaves transiently expressing HaFLS2-1 following treatment with 200 nM flg22^Pst^, flg22^Xcc^, flg22^Xoo^, flg22^Agro^ or flg22^Rso^. Luminescence was monitored for 60 min after peptide addition using the luminol-based assay described in Methods. Curves show mean ± s.d. (*n* = 6 leaf discs per treatment). RLU, relative light units.

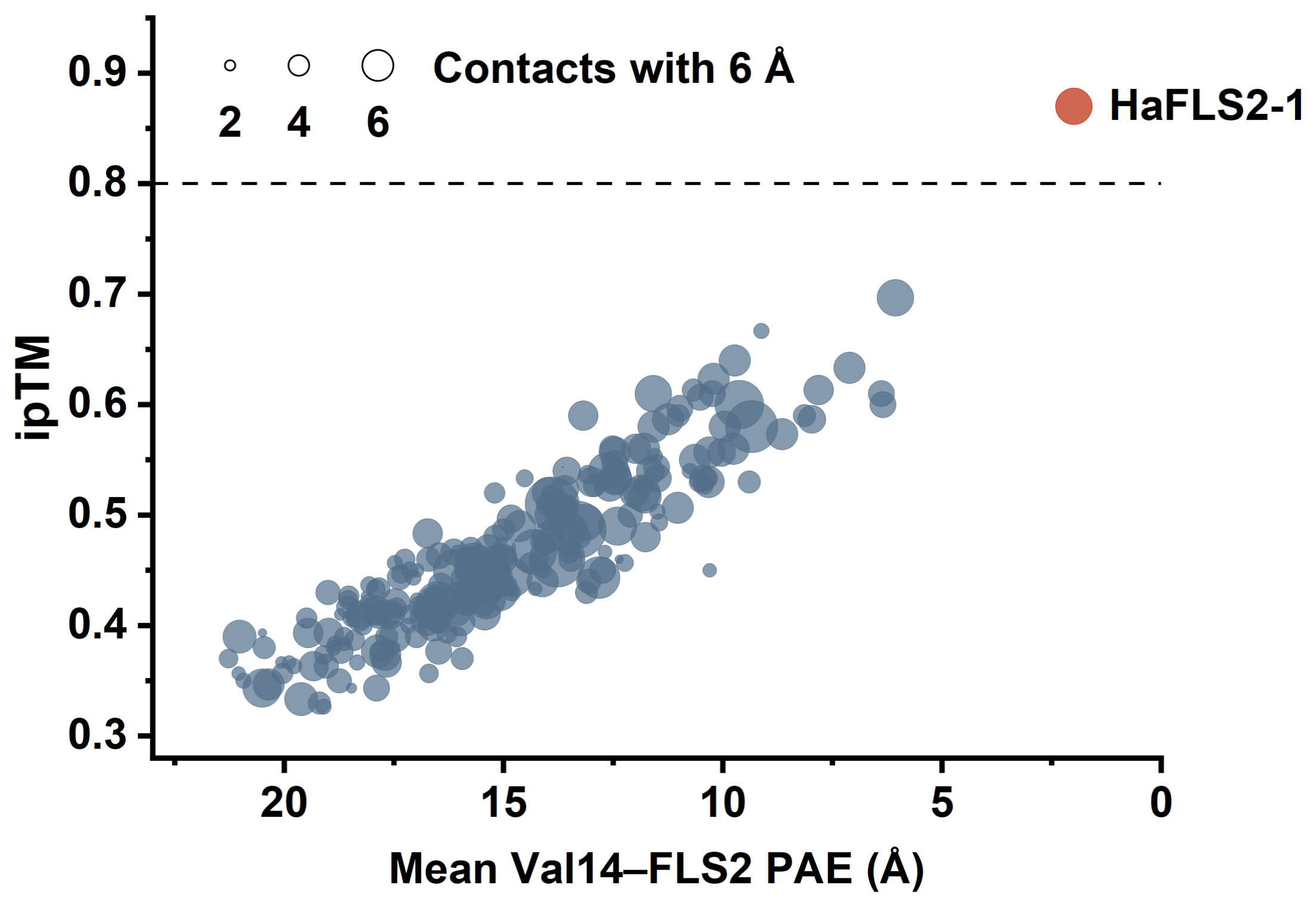

**Figure S4 | Parallel AF3 screening of natural FLS2 homologues with flg22^Xoo^.** AlphaFold 3 (AF3) screening of 256 plant FLS2 homologues modelled in binary complexes with flg22^Xoo^. Each point represents one FLS2 homologue. Mean interface predicted TM score (ipTM) was calculated across three independent AF3 predictions generated with different random seeds. For each prediction, Val14–FLS2 contact predicted aligned error (PAE) was calculated as the mean bidirectional PAE between flg22^Xoo^ Val14 and contacting FLS2 residues; mean PAE was calculated across predictions with defined contacts, as described in Methods. Point size represents the mean number of Val14-contacting FLS2 residues; the contact-distance key is indicated in the plot. HaFLS2-1 is highlighted in red. The dashed line marks ipTM = 0.8. Source values are provided in Supplementary Table S1C.

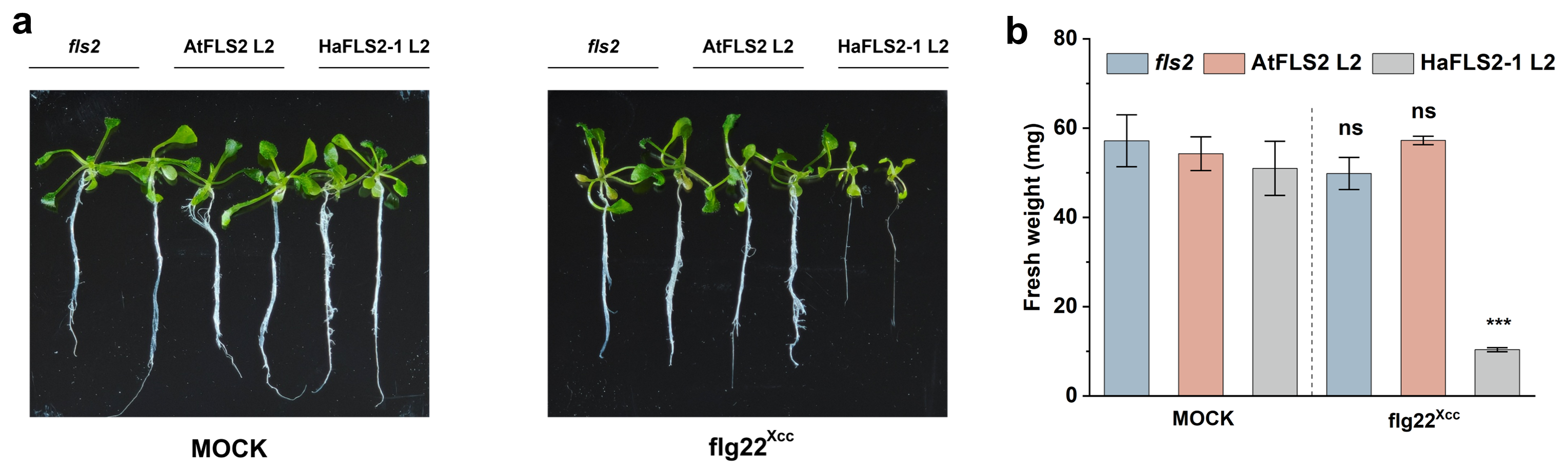

**Figure S5 | flg22^Xcc^-induced growth inhibition in independent *Arabidopsis* transgenic lines. a,** Representative seedlings of the *Arabidopsis thaliana* *fls2* mutant and independent AtFLS2 L2 and HaFLS2-1 L2 transgenic lines following mock or 10 μM flg22^Xcc^ treatment. Seedlings were grown on solid medium for 7 d and then transferred to liquid medium containing flg22^Xcc^ or a DMSO-matched mock medium without peptide for a further 12 d. **b,** Fresh weight of seedlings after treatment. Bars show mean ± s.d. (*n* = 3 seedlings per genotype and treatment). Mock and flg22^Xcc^ treatments were compared within each genotype using two-tailed unpaired *t*-tests, with Welch’s correction for unequal variances. ns, not significant; ****P* < 0.001. L2 denotes an independent transgenic line, complementary to the L1 results in Fig. 1i,j.

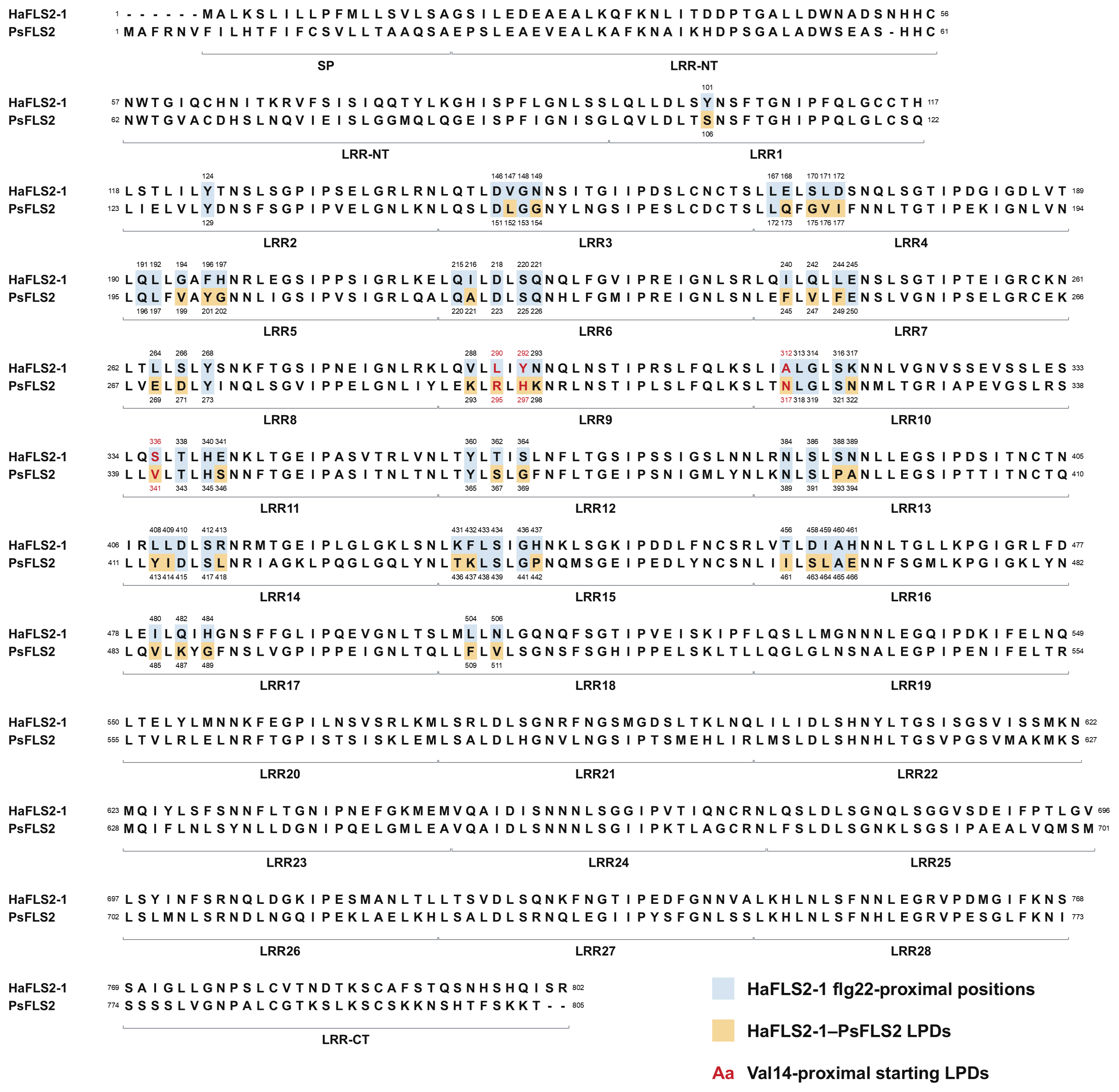

**Figure S6 | HaFLS2-1–PsFLS2 ectodomain alignment defining ligand-proximal difference positions.** The HaFLS2-1 and *Populus simonii* FLS2 (PsFLS2) ectodomain input sequences, including the N-terminal signal peptides, were aligned with LRR boundaries indicated below. Numbering follows each receptor sequence; hyphens denote alignment gaps. Pale blue shading marks HaFLS2-1 positions identified within 6 Å of flg22^Xcc^ across AF3-predicted complexes, together with identical residues at the corresponding PsFLS2 positions. Ochre shading marks ligand-proximal difference positions (LPDs), where PsFLS2 differs from HaFLS2-1 within this ligand-proximal set. Of the 43 LPDs annotated in the alignment, 41 were included in the iterative search; F509L and V511N were not included. The four Val14-proximal starting substitutions (R295L, H297Y, N317A and V341S) are highlighted in red. Substitution notation gives the PsFLS2 residue, position and HaFLS2-1 target residue. Search trajectories and tested variants are provided in Supplementary Tables S3 and S5.

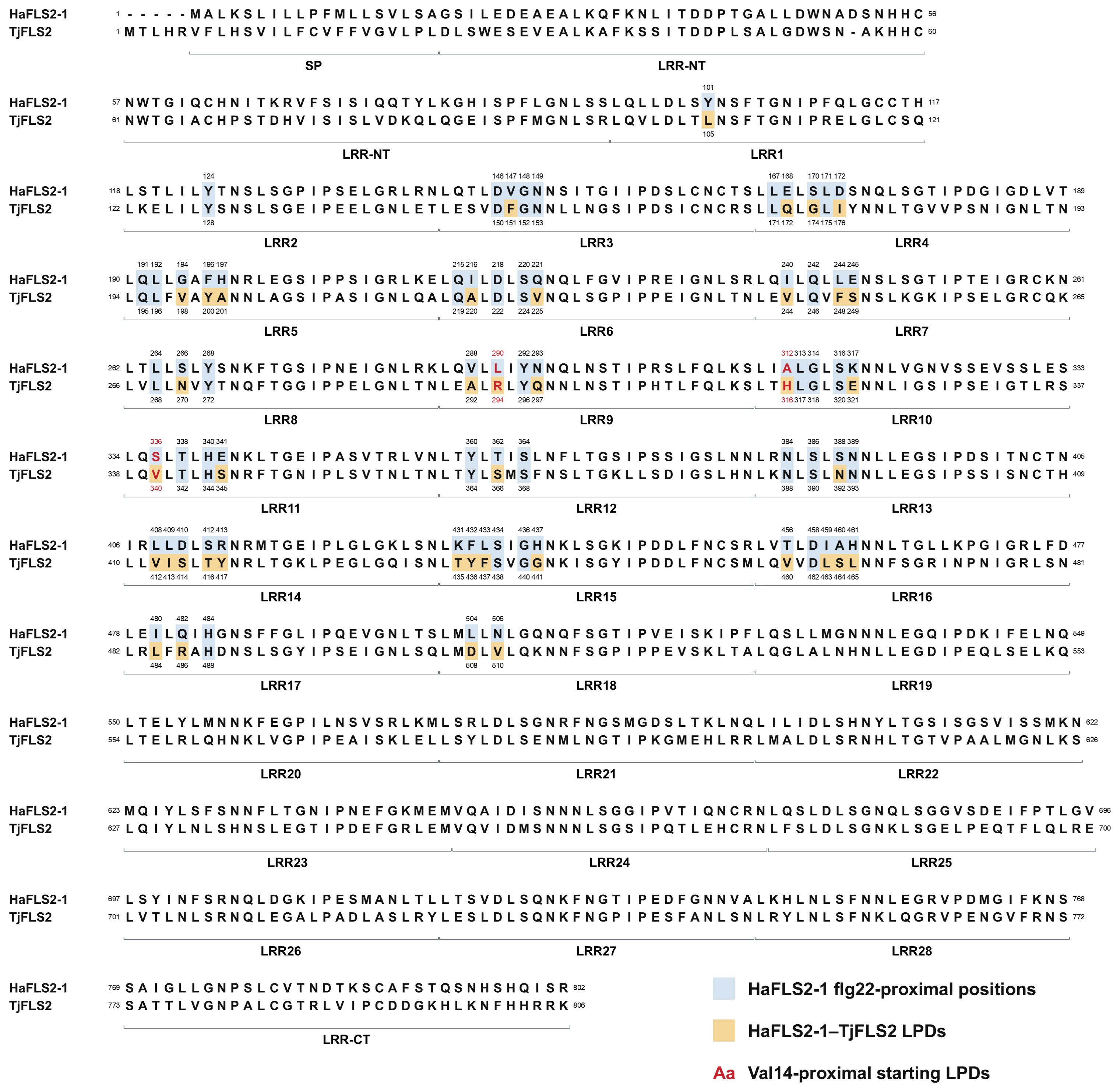

**Figure S7 | HaFLS2-1–TjFLS2 ectodomain alignment defining ligand-proximal difference positions.** The HaFLS2-1 and *Trachelospermum jasminoides* FLS2 (TjFLS2) ectodomain input sequences, including the N-terminal signal peptides, were aligned with LRR boundaries indicated below. Numbering follows each receptor sequence; hyphens denote alignment gaps. Pale blue shading marks HaFLS2-1 positions identified within 6 Å of flg22^Xcc^ across AF3-predicted complexes, together with identical residues at the corresponding TjFLS2 positions. Ochre shading marks the 40 ligand-proximal difference positions (LPDs), where TjFLS2 differs from HaFLS2-1 within this ligand-proximal set. All 40 LPDs were included in the iterative search. The three Val14-proximal starting substitutions (R294L, H316A and V340S) are highlighted in red. Substitution notation gives the TjFLS2 residue, position and HaFLS2-1 target residue. Search trajectories and tested variants are provided in Supplementary Tables S4 and S5.

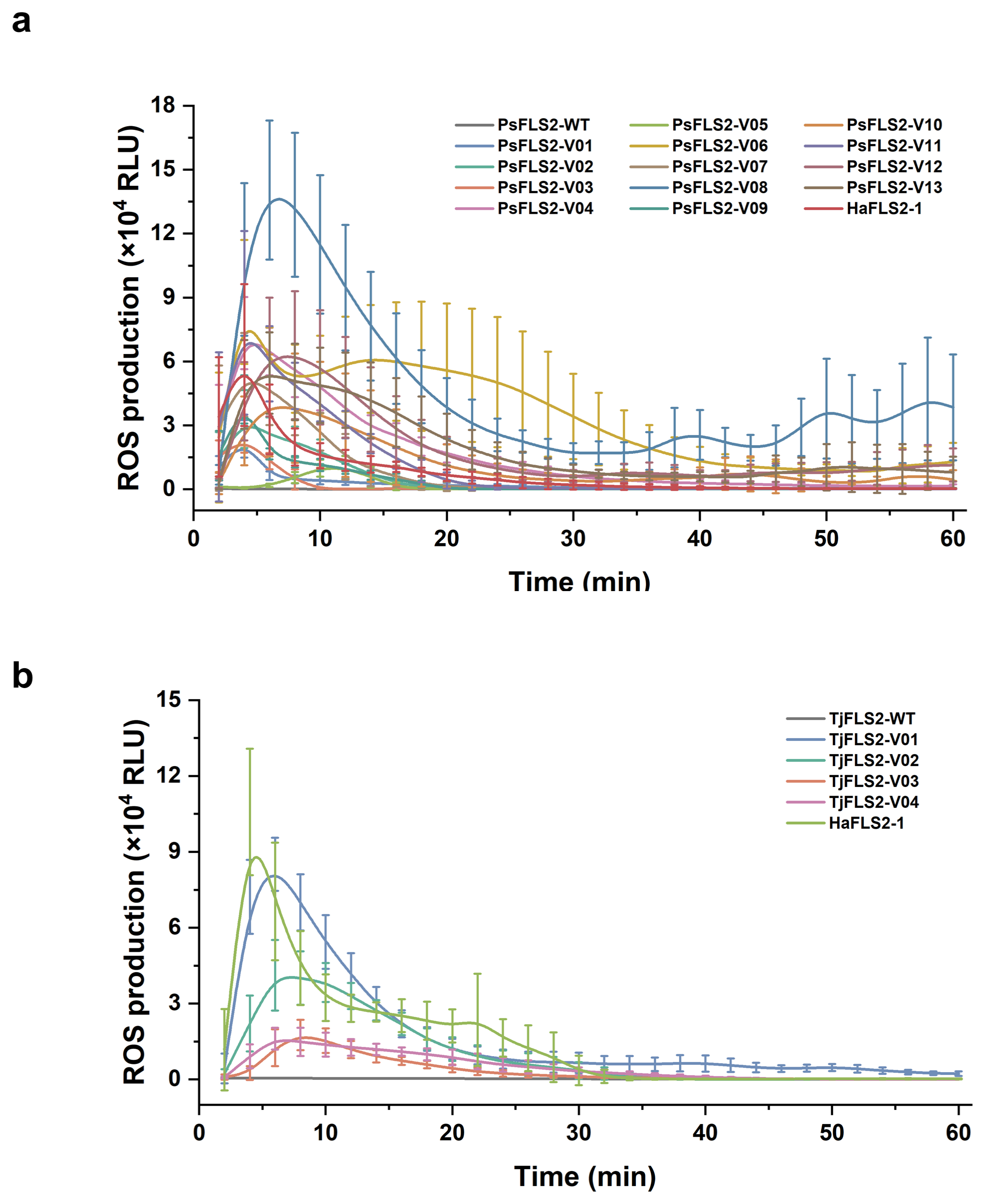

**Figure S8 | ROS kinetics of engineered PsFLS2 and TjFLS2 variants following flg22^Xcc^ treatment. a,** Time-course ROS production in *N. benthamiana* *fls2* leaves transiently expressing wild-type PsFLS2 or the 13 engineered PsFLS2 variants (V01–V13). **b,** Corresponding kinetics for wild-type TjFLS2 and the four engineered TjFLS2 variants (V01–V04). HaFLS2-1 served as the natural positive control in both panels. Leaf discs were treated with 200 nM flg22^Xcc^, and luminescence was monitored for 60 min. Curves show mean ± s.d. (*n* = 8 leaf discs per construct). Variant numbering corresponds to Fig. 2d,g and the experimentally tested constructs in Supplementary Table S5. RLU, relative light units.

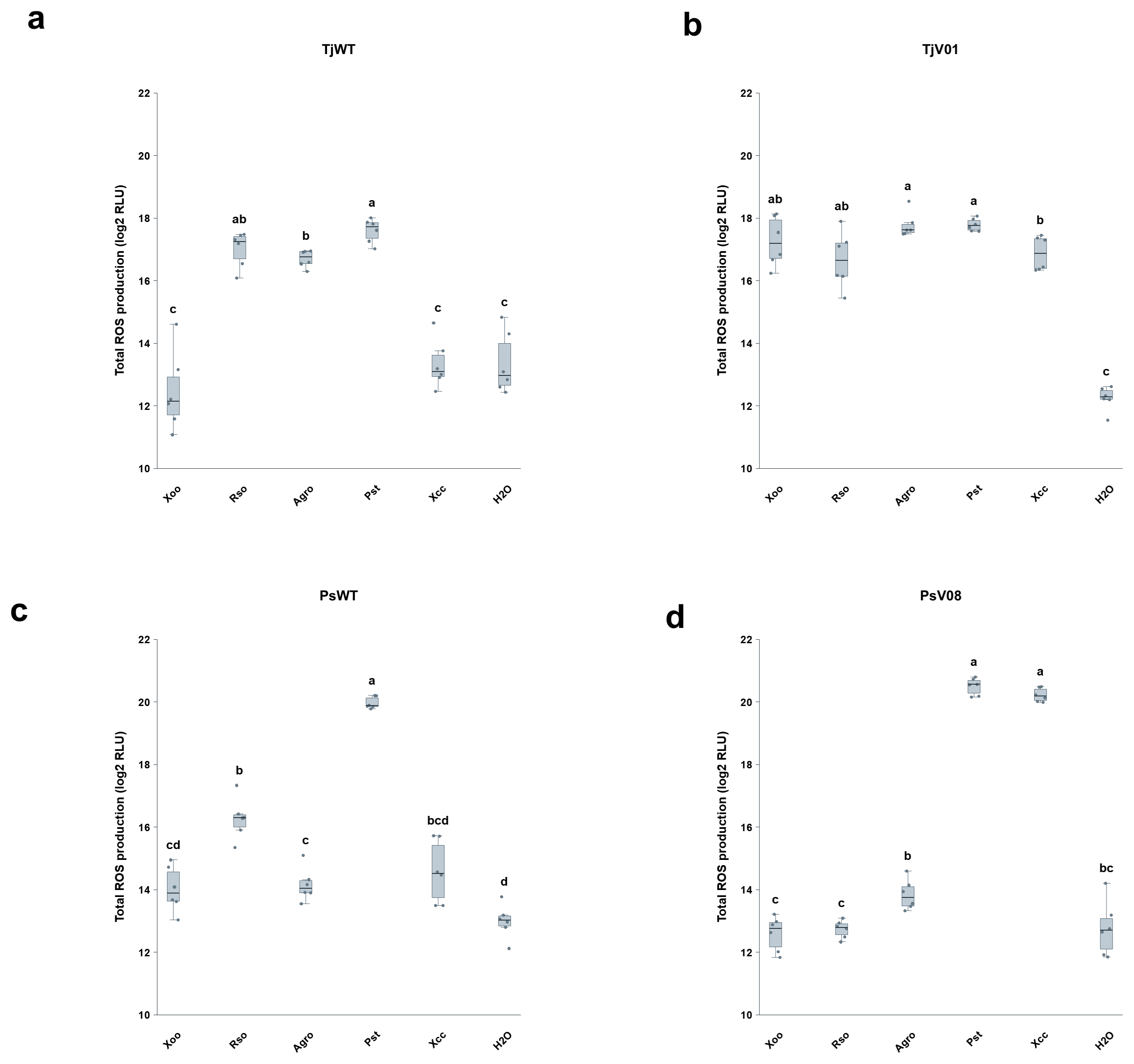

**Figure S9 | Recognition spectra of wild-type and engineered TjFLS2 and PsFLS2 receptors.** Total ROS production in *N. benthamiana* *fls2* leaves transiently expressing wild-type TjFLS2 (a), TjFLS2-V01 (b), wild-type PsFLS2 (c) or PsFLS2-V08 (d), following treatment with 200 nM flg22^Xoo^, flg22^Rso^, flg22^Agro^, flg22^Pst^ or flg22^Xcc^. Water (H_2_O) served as the negative control. Total ROS production over the recording period is shown as log_2_-transformed RLU. Individual measurements are shown as points, with boxes indicating the median and interquartile range (*n* = 6 leaf discs per receptor and treatment). Statistical comparisons were performed separately within each receptor construct using Welch’s one-way ANOVA followed by Games–Howell post hoc tests. Groups sharing no letter differ significantly (*P* < 0.05). Letters are specific to each panel and do not indicate comparisons between receptor constructs. Corresponding TjFLS2-V01 ROS kinetics are shown in Fig. 2i.

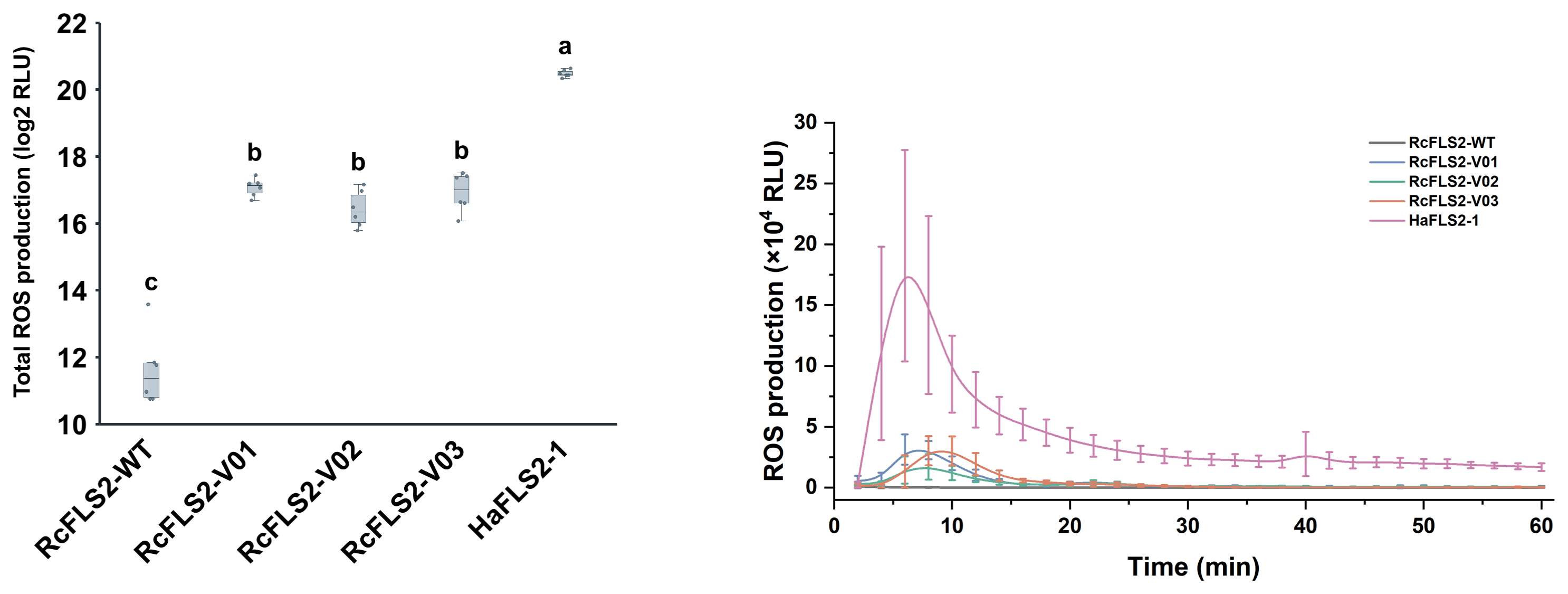

**Figure S10 | flg22^Xcc^-induced ROS production mediated by engineered RcFLS2 variants.** ROS production in *N. benthamiana* *fls2* leaves transiently expressing wild-type *Rosa chinensis* FLS2 (RcFLS2), three engineered RcFLS2 variants (V01–V03) or HaFLS2-1 (positive control), following treatment with 200 nM flg22^Xcc^. Left, total ROS production shown as log_2_-transformed RLU. Individual measurements are shown as points, with boxes indicating the median and interquartile range (*n* = 6 leaf discs per construct). Statistical comparisons used Welch’s one-way ANOVA followed by Games–Howell post hoc tests. Groups sharing no letter differ significantly (*P* < 0.05). Right, corresponding ROS kinetics over 60 min; curves show mean ± s.d. (*n* = 6 leaf discs per construct). Experimentally tested constructs and AF3 search records are provided in Supplementary Tables S5 and S7, respectively.

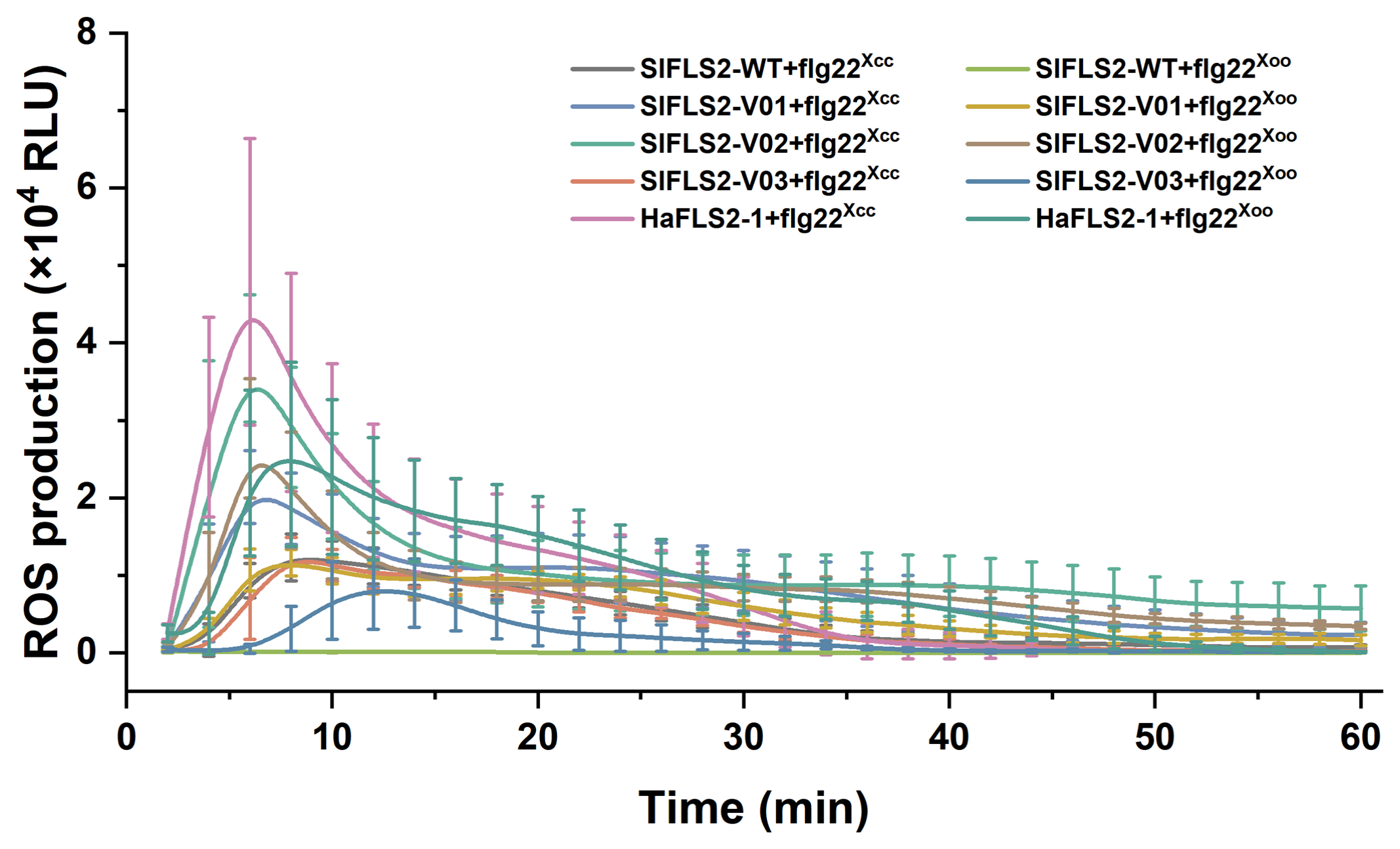

**Figure S11 | ROS kinetics of wild-type and engineered SlFLS2 following flg22^Xcc^ or flg22^Xoo^ treatment.** Time-course ROS production in *N. benthamiana* *fls2* leaves transiently expressing wild-type *Solanum lycopersicum* FLS2 (SlFLS2), the three engineered SlFLS2 variants (V01–V03) or HaFLS2-1 (positive control), following treatment with 200 nM flg22^Xcc^ or flg22^Xoo^. Luminescence was monitored for 60 min. Curves show mean ± s.d. (*n* = 6 leaf discs per receptor and treatment). One representative experiment is shown. The corresponding total ROS measurements and statistical comparisons are shown in Fig. 2l. Variant numbering corresponds to Supplementary Table S5; AF3 search records are provided in Supplementary Table S8. RLU, relative light units.
